# Accelerated and High-Accuracy Variant Calling on Oxford Nanopore Technologies Sequencing Data with the Sentieon DNAscope LongRead and Hybrid Pipelines

**DOI:** 10.1101/2025.11.14.688541

**Authors:** Jinnan Hu, Don Freed, Hanying Feng, Hong Chen, Zhipan Li, Haodong Chen

## Abstract

Oxford Nanopore Technologies (ONT) has emerged as a clinically meaningful long-read sequencing platform, enabled by recent improvements in read accuracy and throughput. However, variant calling on ONT data remains challenging due to platform-specific error profiles, particularly indels in homopolymer regions. Here, we introduce description and comprehensive benchmarks of the Sentieon DNAscope LongRead and DNAscope Hybrid pipelines on ONT whole-genome datasets, demonstrating high accuracy and substantially reduced computational requirements across SNP, Indel, and Structural Variant detection.

Benchmarked against multiple truth sets including GIAB v4.2.1, CMRG, and T2T-Q100, DNAscope LongRead reduced SNP errors by ∼50% relative to Clair3 across five reference samples and showed consistently higher accuracy in all non-long-homopolymer stratifications. Integration of ONT and Illumina short reads via DNAscope Hybrid further improved variant detection: with low depth ONT + Illumina datasets, the pipeline achieved F1 scores of 0.9992 (SNPs) and 0.9979 (Indels), outperforming alternative pipelines. In challenging benchmarks such as T2T-Q100 and CMRG genes, DS-Hybrid reduced SNP and Indel errors significantly, compared to next-best methods. For structural variants, DNAscope pipelines outperformed alternative (e.g. Sniffles2), highlighting the advantages of haplotype-resolved SV detection method.

Despite its accuracy, the Sentieon software suite remains computationally efficient: complete ONT long read FASTQ-to-VCF analysis completed within ∼190 minutes on a 120-vCPU Azure instance, at costs less than $5 USD.

Overall, the DNAscope LongRead and Hybrid pipelines deliver fast, accurate, and scalable germline variant calling for ONT datasets, providing a practical solution for comprehensive whole-genome analysis in both research and clinical genomics.

## Introduction

Over the past decade, next- and third-generation sequencing (NGS and TGS) have become foundational in genomics research and medicine, driven by improvements in read length, accuracy, and accessibility^1^. While short-read sequencing technologies (e.g., Illumina, Element Biosciences, MGI) excel at profiling SNPs and small indels across most of the genome, they remain limited in difficult-to-map regions and in detecting larger or complex structural variation (SV)^2^. Long-read sequencing platforms — particularly Oxford Nanopore Technologies (ONT)—addresses many of these limitations through extended read lengths that enable more confident detection and refinement of SVs and other challenging variants^3,4^. However, ONT error rate, particularly indels, can complicate variant calling and require high coverage along with substantial computational resources^5^.

Recent advances have established Oxford Nanopore Technologies (ONT) as both a high-accuracy and clinically viable long-read sequencing platform. The introduction of ONT’s Q20+ chemistry and R10.4 flow cells has markedly improved base-calling accuracy, with average read approaching Illumina-level performance^6,7^. These improvements have translated directly into clinical utility: ultrarapid ONT whole-genome sequencing in critically ill pediatric patients achieved a median turnaround of just 5.3 days, substantially expediting diagnosis and treatment decisions^8^. Moreover, ONT-based long-read WGS has been shown to complement short-read pipelines by resolving complex structural variants and providing orthogonal confirmation of clinically relevant mutations^9^. Collectively, these studies highlight that ONT sequencing now delivers both the accuracy and operational speed required for comprehensive, time-sensitive genomic analysis in clinical settings.

The benchmark improvements further highlight the value of advanced long-read technologies. The Genome in a Bottle (GIAB) Consortium has progressively expanded high-confidence regions in GRCh38 through incorporation of linked- and long-read data, including more difficult genomic loci^10^. Additionally, draft assembly-based benchmarks derived from high-quality T2T datasets^11,12^ have substantially increased confident small variants and SV calls^13^ in regions previously inaccessible by short reads alone, reinforcing the importance of long reads-enabled characterization of challenging genomic areas.

Building on these previous developments, we developed and reported the DNAscope LongRead (DS-LR) ^14^ and DNAscope Hybrid (DS-Hybrid)^15^ pipelines, which respectively analyze long reads alone or integrate long- and short-read data from the same sample. However, earlier publications did not include Oxford Nanopore Technologies (ONT) datasets or performance evaluations; this study fills that gap by providing comprehensive ONT benchmarks. Compared with existing methods such as Clair3^16^ and Sniffles2^17^, the DNAscope pipelines deliver higher variant-calling accuracy while requiring substantially fewer computational resources.

## Results

### DNAscope LongRead Pipeline Overview

In this study, we present the current version of the DNAscope LongRead pipeline with an optimized ONT model, designed to achieve highly accurate detection of both small and structural variants from Oxford Nanopore sequencing data. As illustrated in Figure 1, DNAscope LongRead accepts FASTQ or BAM inputs and produces VCF outputs containing SNPs, indels, and structural variants. The pipeline is compatible with whole-genome long-read sequencing as well as targeted sequencing applications. Its demonstrated accuracy and flexibility make DNAscope LongRead a robust and practical tool for clinical genomics, particularly in contexts that demand comprehensive and reliable variant detection.

**Figure 1:**
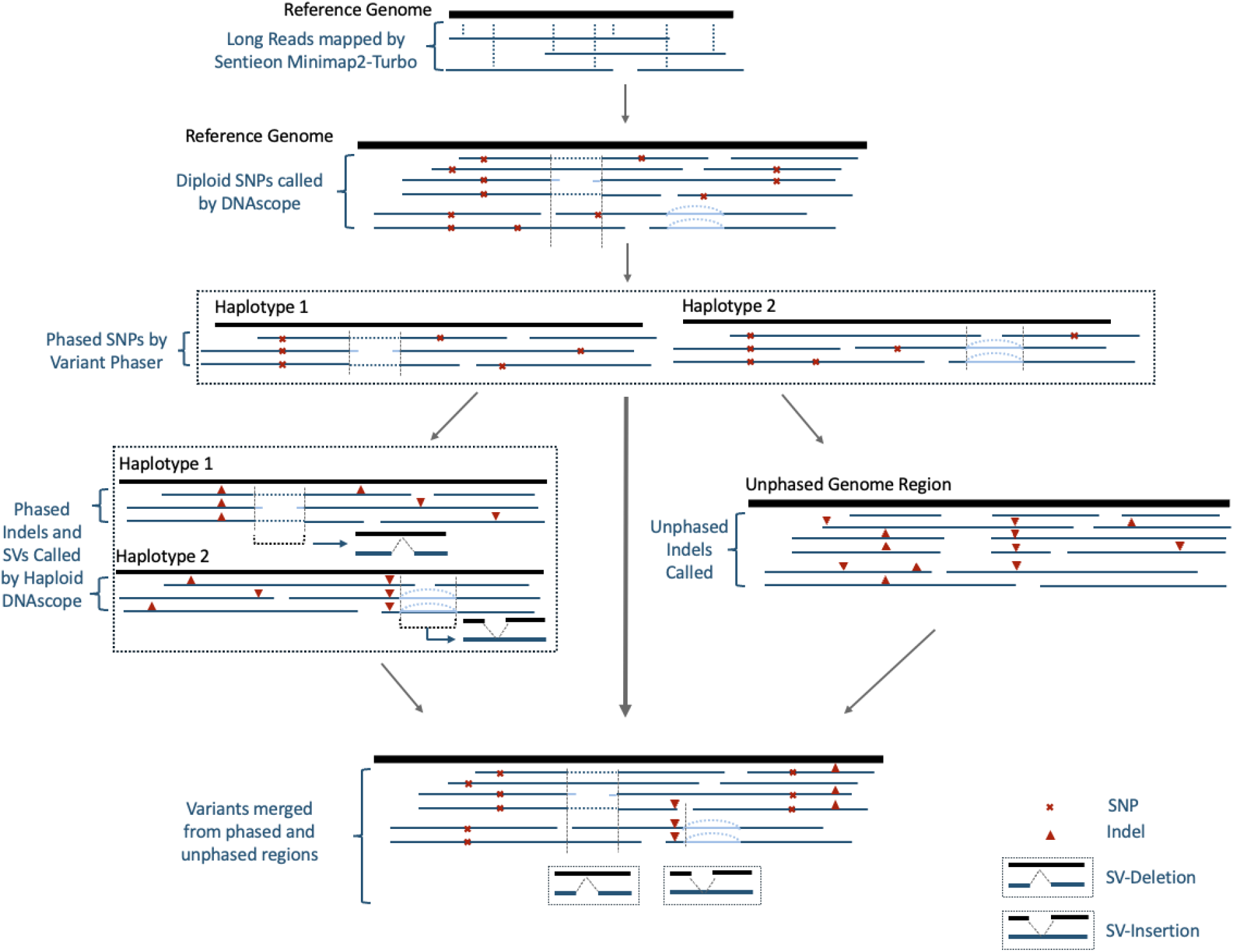
Overview processing steps of the DNAscope LongRead variant calling pipeline.

To evaluate the performance of the DNAscope LongRead and Hybrid pipeline on ONT dataset, we benchmark the pipeline output using a variety benchmarks. We benchmark the small variant (SNP and Indel) VCF using the GIAB v4.2.1, CMRG^18^, and T2T-Q100 benchmarks. SVs identified by the pipeline are assessed using the CMRG and T2T-Q100 SV benchmarks. The runtime of the DNAscope both pipeline is assessed by running the pipelines using a public cloud server.

### Small Variants (SNPs and Indels) from DNAscope LongRead Pipeline

We first evaluated the performance of DNAscope LongRead (DS-LR) on ONT data across five GIAB reference samples (Figure 2, Table S1). The machine learning model used was trained on HG001, HG002, and HG004–HG007, with HG003 held out as an independent test sample. Results were compared against Clair3 (v1.0.8), which serves as the current standard short variant caller in the ONT-recommended analysis pipeline.

**Figure 2.**
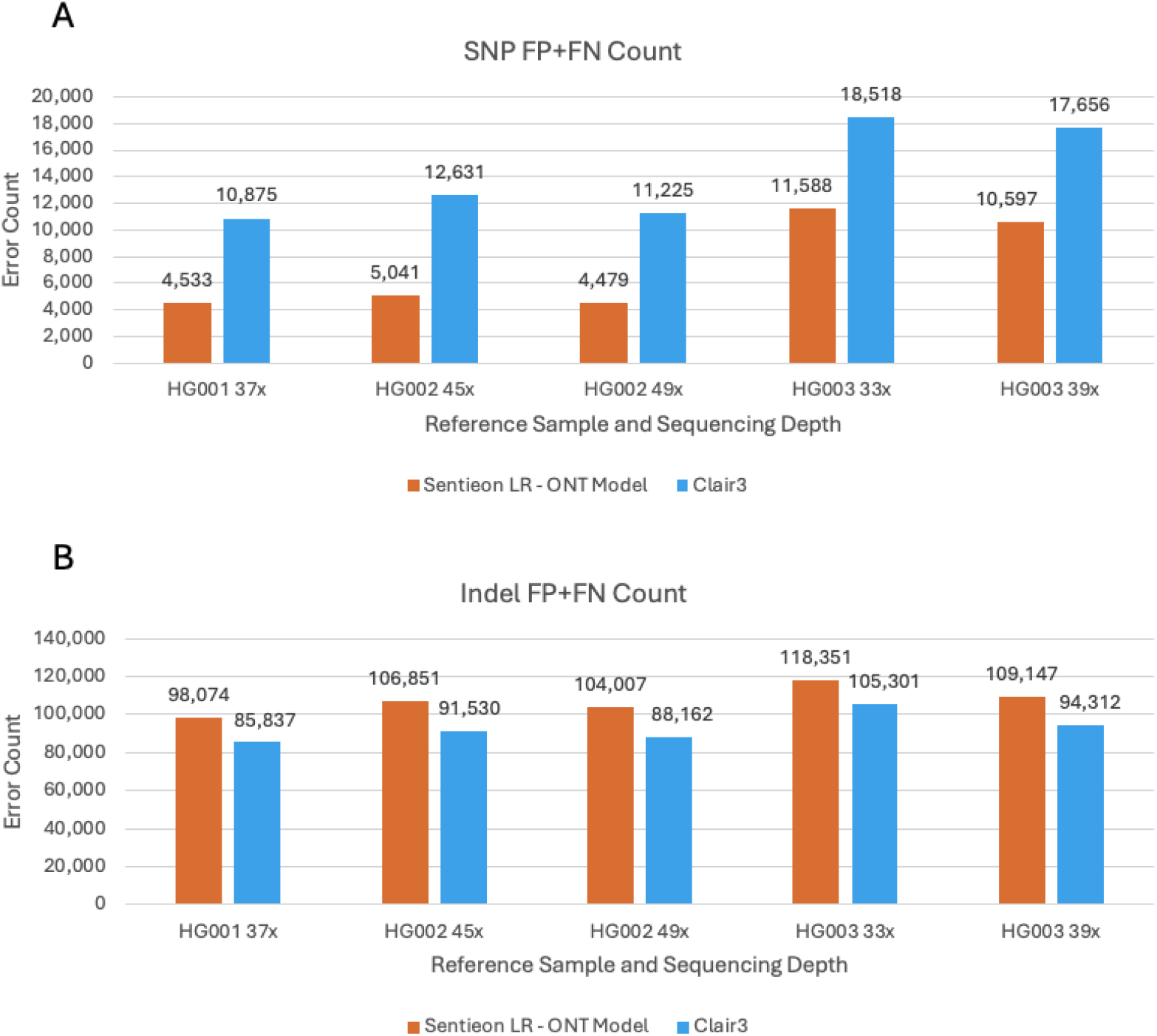
A) SNP and B) Indel error counts across five reference samples at varying sequencing depths.

Due to the intrinsic error profile of ONT reads—particularly insertion and deletion errors within homopolymer regions—Indel detection is generally more challenging than SNP or structural variant detection. As shown in Figure 2, the Sentieon DS-LR pipeline with the ONT-optimized model reduced SNP errors by approximately 50% compared to Clair3. As expected, HG003, which was excluded from model training, exhibited a higher error count than HG001 and HG002, and a similar trend was observed for Clair3.

To further characterize error sources, we analyzed SNP and Indel accuracy within different GA4GH stratified genomic contexts^19^, reporting F1 scores (instead of error counts) to normalize for varying region sizes (Table 1, Table S2). As expected, both callers showed the greatest challenges in homopolymer regions longer than 11 bp, where Indel detection was largely lost. Accuracy improved slightly for shorter homopolymers (4–11 bp) but remained lower than in other genomic categories. Outside of homopolymer regions—even within “low-mappability or segmental duplication” areas—DS-LR achieved accuracy levels comparable to other sequencing platforms, peaking in “easy” regions where both SNP and Indel F1 scores approached 0.999. Notably, across all non-long homopolymer (>7 bp) regions, the Sentieon pipeline outperformed Clair3 for both SNP and Indel detection.

**Table 1.**
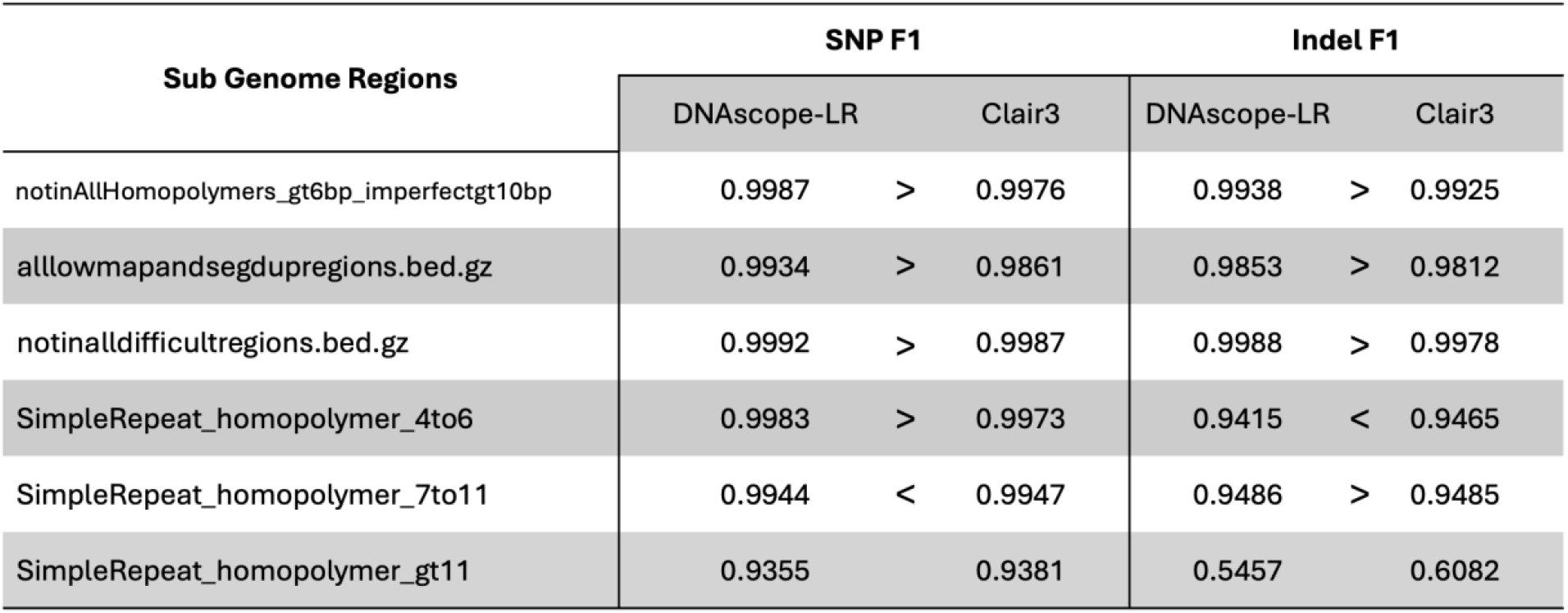
Accuracy of selected stratification genomic regions.

| Sub Genome Regions | SNP F1 |  |  | Indel F1 |  |  |
| --- | --- | --- | --- | --- | --- | --- |
|  | DNAScope-LR |  | Clair3 | DNAScope-LR |  | Clair3 |
| notinAllHomopolymers_gt6bp_imperfectgt10bp | 0.9987 | > | 0.9976 | 0.9938 | > | 0.9925 |
| allowmapandsegdupregions.bed.gz | 0.9934 | > | 0.9861 | 0.9853 | > | 0.9812 |
| notinAlldifficultregions.bed.gz | 0.9992 | > | 0.9987 | 0.9988 | > | 0.9978 |
| SimpleRepeat_homopolymer_4to6 | 0.9983 | > | 0.9973 | 0.9415 | < | 0.9465 |
| SimpleRepeat_homopolymer_7to11 | 0.9944 | < | 0.9947 | 0.9486 | > | 0.9485 |
| SimpleRepeat_homopolymer_gt11 | 0.9355 |  | 0.9381 | 0.5457 |  | 0.6082 |

### Small Variants from DNAscope Hybrid Pipeline

Next, to evaluate the accuracy of the DNAscope Hybrid (DS-Hybrid) pipelines at varying depths, we used a HG002 ONT dataset down sampled to depths of 5x, 7.5x, 10x, 15x, 20x, and 30x, paired with 35x Illumina (ILMN) short-read data. Additionally, we also analyzed each depth of ONT datasets independently without short reads using the DS-LR pipeline. Further, we included other datasets for comparison: Illumina (35x) data analyzed with DNAscope (short reads with linear reference genome)^20^, and ONT (30x) analyzed by Clair3 v1.0.8 (Clair3). It should be noted DS-LR accuracy demonstrated in this section was from an older version compared to the above DS-LR dedicated benchmark, and thus accuracy was also lower.

We initially investigated genome-wide accuracy using the NIST v4.2.1 benchmark (Figure 3 A-B, Table S3). This analysis demonstrated that higher depths of long-read sequencing yield greater accuracy, with the highest accuracy observed in the combined 30x ONT + 35x ILMN datasets. Further, Hybrid Indel accuracy is higher than the other evaluated methods, even when using only 5x coverage for long reads. 10x ONT + 35x Illumina seems to have a good balance between the cost of reagents and results. At this coverage level the pipeline has 2,180 Indel errors and 5,662 SNP errors when evaluated on the GIAB v4.2.1 benchmark, for an F1 of 0.9979 and 0.9992 respectively.

**Figure 3.**
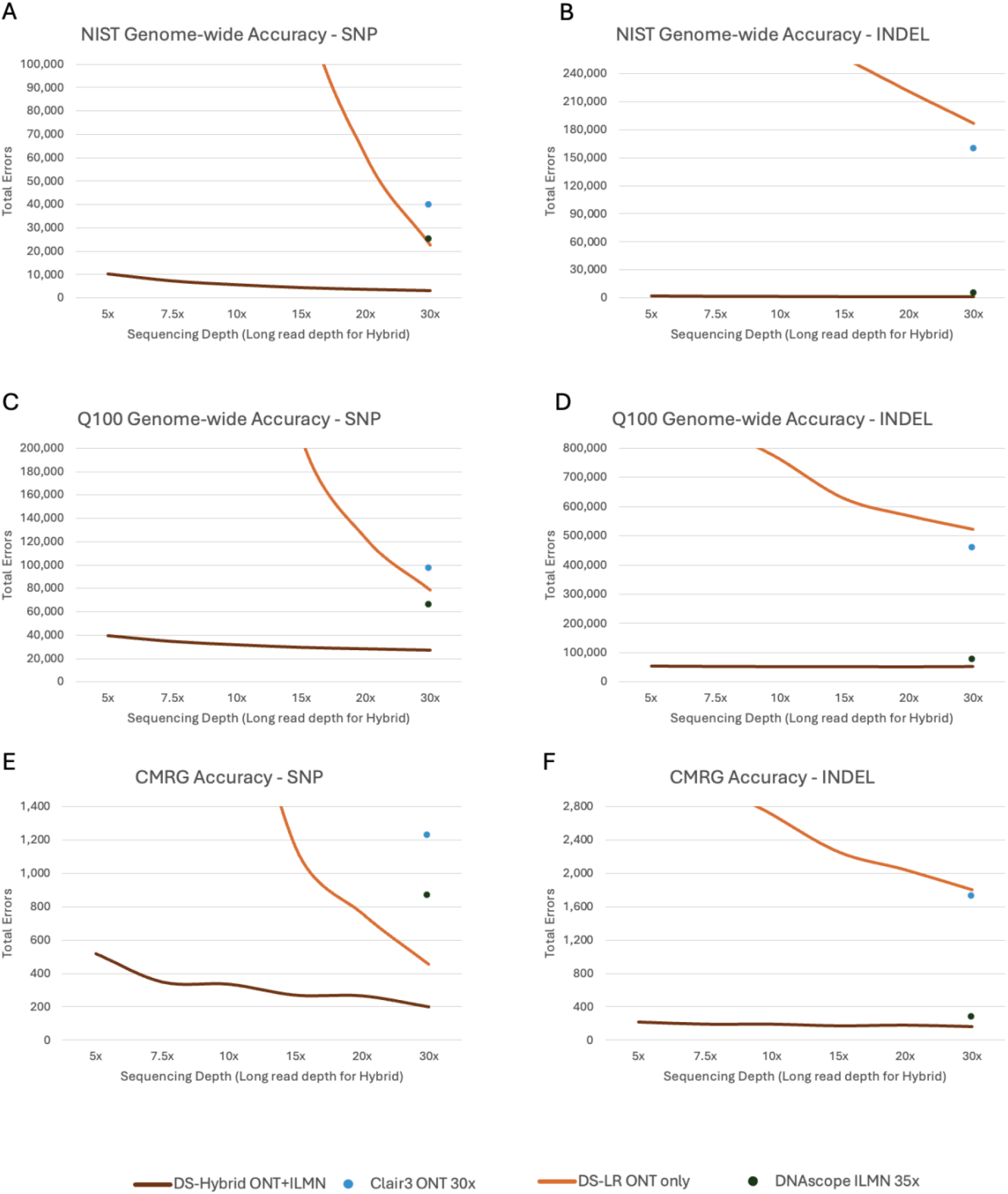
Genome-wide Accuracy - Total Errors of A) SNP and B) Indel in GIAB v4.2.1; C) SNP and D) Indel in T2T-Q100; E) SNP and F) Indel in challenging medically relevant gene (CMRG) regions. DS-Hybrid ONT+ILMN and DS-LR ONT only are shown with curves covering 5x-30x long reads depths. Clair3 ONT and DNAscope ILMN are shown at full depth.

Total errors are much higher with the T2T-Q100 benchmark than the v4.2.1 benchmarks, as it contains more challenging regions (Figure 3 C-D, Table S4), making it more suitable for benchmarking new high accuracy variant callers. In the T2T-Q100 benchmark, the DS-hybrid pipeline has fewer errors relative to short or long reads alone pipelines for both SNPs and Indels. Comparing with the hybrid pipeline at 10x long-read coverage, SNP errors are reduced by 51% relative to the next-best pipeline (DNAscope ILMN) and Indel errors are reduced by 27% as well.

CMRG regions, which encompass 273 medically relevant genes, demonstrated substantial benefits from hybrid short and long-read data. Long reads alone cannot capture each variant correctly, while the Hybrid pipeline still showed its improved accuracy (Figure 3 E-F, Table S5). The improved accuracy will likely lead to an improved diagnostic rate and other clinical utility.

While the DS-Hybrid pipeline has excellent performance on HG002, we wanted to further assess the performance on additional datasets to ensure that the approach used by the pipeline extends to other samples (Figure 4, Table S6). We then applied the pipeline to two additional GIAB samples. The results, measured as SNP and Indel combined total errors (FP+FN), were compared to three Illumina short reads pipelines (Dragen and DNAscope PanGenome^21^ and only with standard linear genome). ONT only results were not shown here since they are much higher than hybrid or short reads only accuracy. Figure 4 shows DS-Hybrid call sets are consistently more accurate than other three short reads call sets among the tested 3 samples, highlighting its robustness and adaptability. Pangenome information could also be integrated into DS-Hybrid pipeline in future and thus provide even higher accuracy.

**Figure 4.**
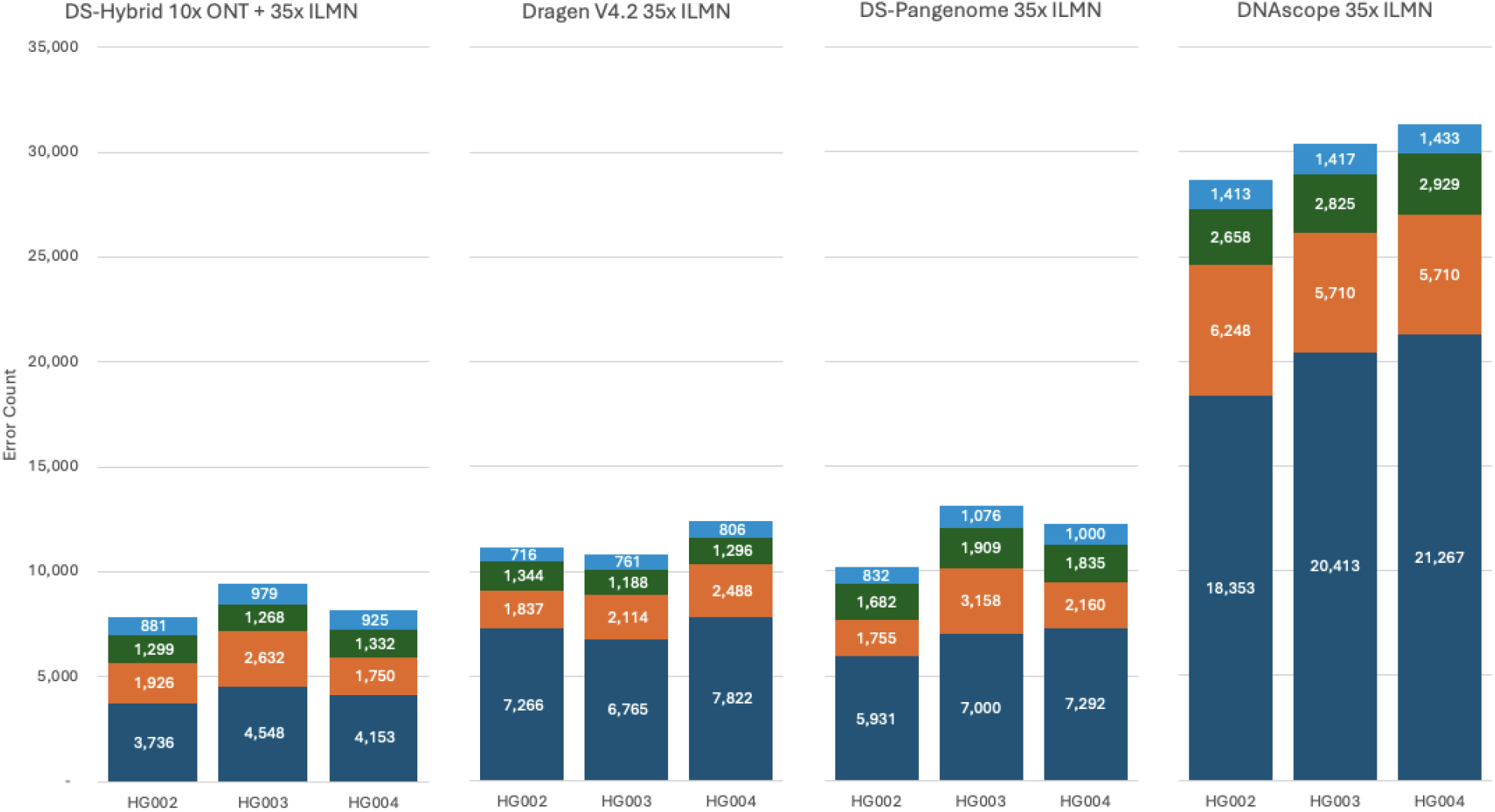
SNP/Indel accuracy over HG002-004 reference samples. False positive and false negative counts for SNPs and Indels are listed separately.

### Structural Variants (SVs)

Both DS-Hybrid and DS-LR pipelines utilize long reads only to call structural variant (SV), so their accuracy curves were combined. We analyzed down sampled long read datasets and evaluated SV calling accuracy using the T2T-Q100 or CMRG SV truth. DNAscope (short reads on linear genome) analyzing 35x ILMN dataset were included as comparison.

While short-read pipelines demonstrate high accuracy for SNP and Indel detection, these pipelines, without using pangenome information, have much lower accuracy for SV calling (Figure 6 A-B, Table S7), especially recall. In contrast, long-read pipelines have higher measured SV accuracy, even at lower depths. The DS-Hybrid/LR pipeline outperformed Sniffles2 in this SV benchmark. To further validate pipeline performance in clinical setting, we assessed SV accuracy using the CMRG SV benchmark (Figure 6 C-D, Table S8). Performance on the CMRG SV benchmark was consistent with the performance observed in the larger draft Q100 SV benchmark, underscoring the advantage of long-read sequencing in SV detection.

**Figure 6.**
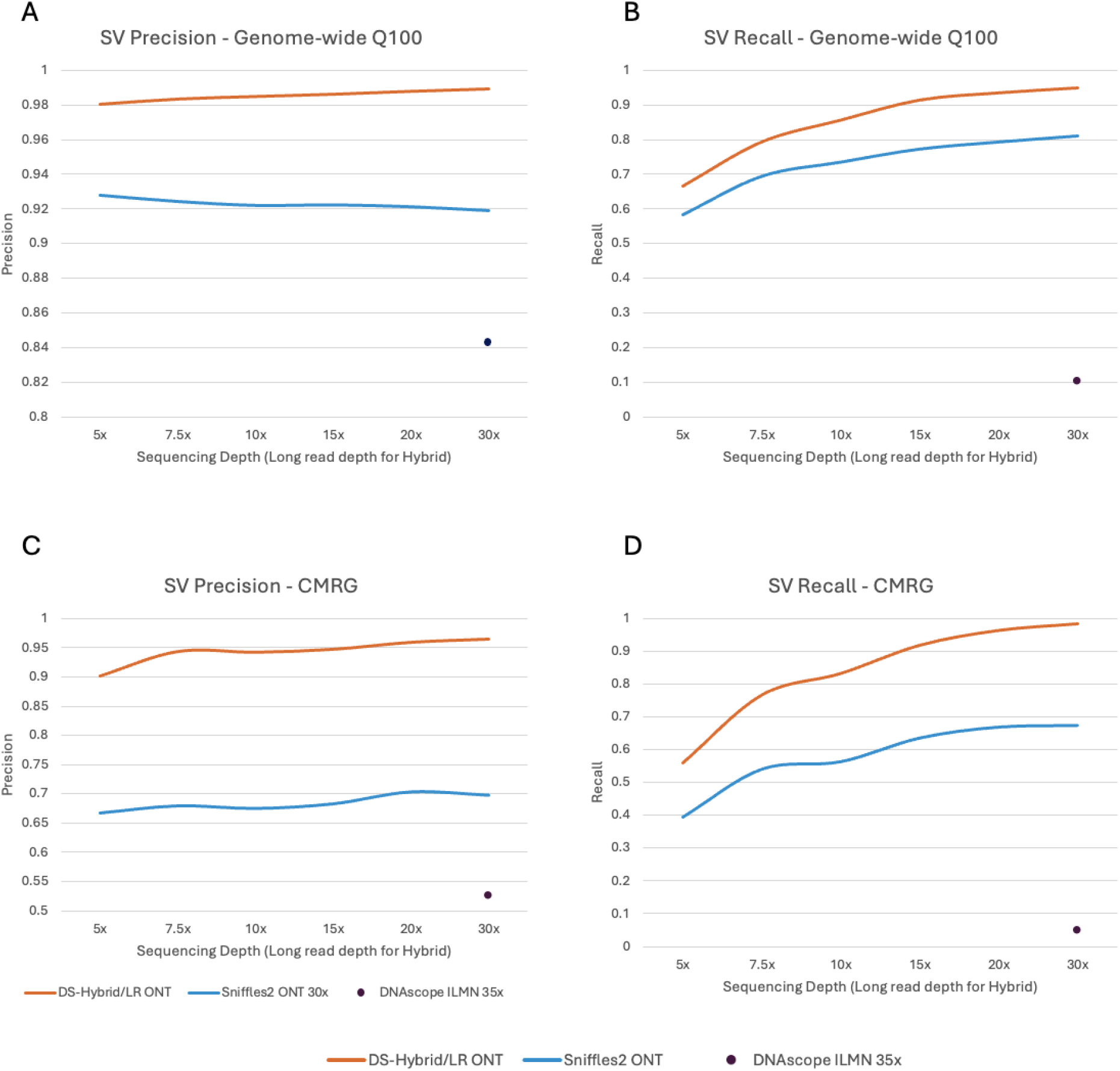
Structural variation (SV) accuracy as measured by A) precision and B) Recall on the T2T-Q100 benchmark; C) Precision and D) Recall on the CMRG SV benchmark. DS-Hybrid/LR ONT and Sniffles2 ONT are shown as curves covering 5x-30x long-read depths. DNAscope ILMN is shown at full depth accuracy.

### Copy Number Variation (CNV)

The DS-Hybrid pipeline also outputs CNV results using the CNVscope module. The results are identical to those reported in our previous study with PacBio HiFi + Illumina data^15^ and are therefore not repeated here, as CNVscope relies exclusively on short-read input.

### Compute Resource Benchmark

A major challenge in whole genome sequencing secondary analysis is the long runtime, high cost of compute, and requirements for specialized hardware for obtaining an adequate TAT. The Sentieon software addresses these issues by running efficiently on commodity (x86 or Arm) CPU servers or workstations, offering accelerated runtimes, improved consistency, and high accuracy compared to other tools.

To assess the runtime of the Sentieon software, we tested three Sentieon pipelines – the DS-Hybrid pipeline with 10x ONT HiFi and 35x Illumina data, DS-LR with 30x ONT data, and DNAscope with 30x Illumina data. The benchmark assessed the runtime performance of alignment, preprocessing and SNP/Indel/SV/CNV calling. A 120 thread Azure instance (Standard HG120rs V3) was used as computation environment. The results for runtime, core-hours, and compute cost are shown in Table 2 below. The DNAscope LongRead and DNAscope pipeline runtime was previously published^22^.

**Table 2.** Compute resource benchmark for DNAscope pipelines. Benchmark environment is Azure Standard HB120rs v3 (120 vCPUs, 456 GiB memory, 512GB premium SSD), runtime and on-demand compute cost are displayed. The DNAscope Hybrid pipeline outputs SNP/Indel/SV/CNV, the DNAscope LongRead pipeline outputs SNP/Indel/SV, DNAscope short reads outputs SNP/Indel/CNV.

| Pipeline | DNAscope Hybrid | DNAscope LongRead | DNAscope (short reads) |
| --- | --- | --- | --- |
| Dataset | 10x ONT + 35x ILMN | 30x ONT | 30x ILMN |
| Alignment (min) | 21.8 | 32.9 | 9.7 |
| Preprocessing (min) | 2.0 | 0.0 | 1.4 |
| Variant Calling (min) | 79.9 | 55.4 | 7.8 |
| Total Runtime (min) | 103.7 | 88.3 | 18.9 |
| Core-hours | 207.4 | 176.6 | 37.8 |
| On-demand (\$) | 6.2 | 5.3 | 1.1 |
| Spot (\$) | 0.62 | 0.53 | 0.11 |

The DNAscope Hybrid pipeline is actively being developed, with future releases expected to show incremental improvements in computational efficiency and accuracy. Benchmarking results indicate that all three Sentieon pipelines completed the fastq-to-VCF analysis in approximately 20 minutes to less than 105 minutes for a cost of between $0.11 and $6.20 depending on data and spot or on-demand pricing.

### Datasets and Methods

### Datasets used in this study

FASTQ files used in this study were downloaded from publicly available sources:

- ONT Dataset:

gs://deepvariant/ont-case-study-testdata/HG002_R104_sup_merged.50x.bam;

gs://deepvariant/ont-case-study-testdata/HG003_R1041_Guppy6_sup_2_GRCh38.pass.bam;

gs://deepvariant/ont-case-study-testdata/HG003_R104_sup_merged.80x.bam;

gs://deepvariant/ont-case-study-testdata/HG004_R104_sup_merged.40x.bam

Illumina: pFDA Truth Challenge V2^23^, HG002+HG003+HG004 Download link: https://precision.fda.gov/challenges/10/intro

Benchmark small variant (SNV and indel) VCF files were downloaded from publicly available sources:

- GIAB v4.2.1 SNP/Indel: https://ftp-trace.ncbi.nlm.nih.gov/ReferenceSamples/giab/release/AshkenazimTrio/
- T2T-Q100 v0.019: https://ftp-trace.ncbi.nlm.nih.gov/ReferenceSamples/giab/data/AshkenazimTrio/analysis/NIST_HG002_DraftBenchmark_defrabbV0.019-20241113/
- CMRG v1.00: https://ftp-trace.ncbi.nlm.nih.gov/ReferenceSamples/giab/release/AshkenazimTrio/HG002_NA24385_son/CMRG_v1.00/GRCh38/SmallVariant/

Benchmark SV VCF files were downloaded from publicly available sources:

- T2T-Q100 v0.019 SV: https://ftp-trace.ncbi.nlm.nih.gov/ReferenceSamples/giab/data/AshkenazimTrio/analysis/NIST_HG002_DraftBenchmark_defrabbV0.019-20241113/
- CMRG v1.00: https://ftp-trace.ncbi.nlm.nih.gov/ReferenceSamples/giab/release/AshkenazimTrio/HG002_NA24385_son/CMRG_v1.00/GRCh38/StructuralVariant/

### DNAscope LongRead Pipeline

The DNAscope LongRead pipeline^14^ was first introduced in collaboration with PacBio in 2022, and subsequent releases have extended support to Oxford Nanopore Technologies (ONT) data, further improving performance and versatility. The pipeline accepts FASTQ or BAM files and outputs SNP, Indel, and SV calls in VCF format, applicable to human whole-genome and targeted sequencing assays.

As illustrated in Figure 1, the current version employs Sentieon Minimap2-Turbo for long-read alignment and DNAscope for diploid SNP calling. Identified SNPs are used by the Variant Phaser module to generate two haplotype sequences, enabling accurate detection of haploid indels and structural variants (SVs were called by DNAscope LongReadSV module); indels from unphased regions are also retained. A pre-trained machine-learning model is applied to filter and correct platform-specific error patterns before merging all variant types into the final output.

DNAscope LongReadSV module is integrated in the DNAscope LongRead and Hybrid pipeline. It performs haplotype-resolved SV calling and works well with both PacBio HiFi and ONT data, supporting various sequencing chemistry versions and base callers.

The DNAscope LongRead pipeline is implemented within the Sentieon software suite—a highly optimized commercial toolkit for biological data processing. Multiple internal modules are orchestrated through the “sentieon-cli” command-line interface, which enables users to execute the complete workflow by specifying input files, output paths, and key parameters.

### DNAscope Hybrid Pipeline

As been described in the publication^15^, The Sentieon DNAscope Hybrid pipeline integrates short- and long-read sequencing data from the same sample to achieve comprehensive and highly accurate variant detection. The pipeline accepts FASTQ or BAM files and outputs SNP, Indel, SV, and CNV calls in VCF format, applicable to human whole-genome and targeted sequencing assays.

Short reads are aligned with Sentieon BWA-turbo and long reads with Sentieon Minimap2-turbo. DNAscope Hybrid performs an initial round of variant calling using both data types with sensitive parameters. Regions showing potential discrepancies—identified by the “hybrid_select.py” script based on genotype disagreement or low-confidence mappings—are selected for refinement. In these regions, long-read alignments are split and re-aligned to guide accurate short-read placement, after which a second variant-calling pass is conducted using the updated short-read alignments.

Results from both passes are merged, genotyped, and filtered using a DNAscope machine-learning model trained on combined short- and long-read datasets (e.g., Illumina and ONT), with the HG003 sample held out for validation. The final normalized VCF includes high-confidence SNPs and Indels. Structural variants are identified using the DNAscope LongReadSV algorithm, and CNVs are detected via the short-read-based CNVscope module. The DNAscope Hybird pipeline can also be run via Sentieon CLI.

### Sentieon Alignment Modules

Accelerated implementations of BWA-MEM^24^ and minimap2^25^ were introduced in the Sentieon software package in 2017^26^ and the 2020.10.04 release, respectively. Both tools generate results consistent with their open-source counterparts while achieving approximately 2–3× faster performance. In the 2023.08 release, a new “turbo” mode for both aligners further improved whole-genome sequencing (WGS) alignment runtimes by about 2× through model-based optimization of computational resources. The turbo versions of these aligners were used in all DNAscope pipelines.

### Software Tools Included as Baselines or for Benchmarking

DNAscope (short reads) served as the baseline for SNP and indel calling from short-read data. Its accuracy and runtime have been improved since the original publication, primarily due to updates to the ModelApply module and retrained machine-learning models. A recent enhancement, DNAscope PanGenome (DS-PanGenome)^21^, extends the short-read DNAscope pipeline to graph-based references. By leveraging pangenome graphs, DS-PanGenome improves alignment and variant-calling accuracy in highly polymorphic and structurally complex genomic regions. The accuracy improvements are demonstrated in Figure 4.

The DNAscope SV caller, included in this study as the short-read structural-variant (SV) performance baseline, operates in three stages: (1) detection of abnormal alignments (split, unmapped, or discordant read pairs) followed by local assembly and filtering of low-support haplotypes; (2) identification of consistent haplotype pairs representing candidate SV breakends; and (3) integration of supporting evidence from discordant read pairs to validate candidates and assign quality scores using a machine-learning model.

Clair3 (v1.0.8)^16^ and Sniffles2 (v2.5.3)^17^ were used to generate small and structural variants under default settings from long-read-only datasets. DRAGEN accuracy metrics were obtained from VCF files provided in a recently published benchmarking study^27^.

Benchmarking of SNVs and indels was performed using hap.py with the GIAB v4.2.1, T2T-Q100, and CMRG benchmark VCF and BED files. Structural-variant benchmarking was conducted using hap-eval^28^, an open-source SV evaluation tool developed by the Sentieon team.

## Supplementary Datasheets

Table S1-S8 contains datasheets and more detailed information for figures in the main manuscript.

## Code Availability

Scripts used in this study can be found in https://github.com/Sentieon/sentieon-cli. Sentieon software is a commercial software package, and free trials are available upon request.

## Competing Interests

All authors are employees of Sentieon Inc. and own stock options as part of their standard compensation package. The reported method and pipeline are part of a commercial software package developed and licensed by the company (Sentieon Inc) where the authors are employed.

